# Zebrafish heme oxygenase 1a is necessary for normal development and macrophage migration

**DOI:** 10.1101/2021.04.07.438802

**Authors:** Kaiming Luo, Masahito Ogawa, Anita Ayer, Warwick J Britton, Roland Stocker, Kazu Kikuchi, Stefan H Oehlers

## Abstract

Heme oxygenase function is highly conserved between vertebrates where it plays important roles in normal embryonic development and controls oxidative stress. Expression of the zebrafish heme oxygenase 1 genes are known to be responsive to oxidative stress suggesting a conserved physiological function. Here we generate a knockout allele of zebrafish *hmox1a* and characterize the effects of *hmox1a* and *hmox1b* loss on embryonic development. We find that loss of *hmox1a* or *hmox1b* causes developmental defects in only a minority of embryos, in contrast to *Hmox1* gene deletions in mice that causes loss of most embryos. Using a tail wound inflammation assay we find a conserved role for *hmox1a*, but not *hmox1b*, in normal macrophage migration to the wound site. Together our results indicate zebrafish *hmox1a* has clearly a partitioned role from *hmox1b* that is more consistent with conserved functions of mammalian Heme oxygenase 1.

## Introduction

Heme oxygenases are a highly conserved family of proteins that catalyze the breakdown of heme to carbon monoxide (CO), ferrous iron and biliverdin [1, 2]. In mammals, there are two isoforms of heme oxygenase (HMOX): the inducible HMOX1, and the constitutively expressed HMOX2 [3]. Both isoforms contain a conserved 24-amino acid sequence known as the “heme binding pocket”, within which a conserved histidine residue acts as the ligand for heme iron [4, 5]. Heme is used as substrate and cofactor for both isoforms, but the physiological properties and regulation of the two proteins are different [4].

Under basal conditions in mammals, HMOX1 is highly expressed in tissues such as spleen, liver, and bone marrow, while it is almost undetectable in other tissues [6]. Expression of HMOX1 is induced by multiple stresses including oxidative stress, heme, iron starvation, inflammatory cytokines, and physical tissue injury [7, 8]. Conversely, mammalian HMOX2 is constitutively and highly expressed in the brain and testes and does not generally respond to stress conditions [4, 9, 10].

The role of HMOX1 in regulating the immune response has been demonstrated in models of tissue injury or different diseases such as ischemic lung/liver injury, atherosclerosis, and non-hemochromatosis liver diseases [11-14]. In 1999, the first case of HMOX1 deficiency was reported in a young boy. He showed severe inflammatory phenotypes and accelerated atherosclerosis. The patient died when he was only six years old [15]. This case suggests a critical role of Hmox1 in protecting humans against inflammation and atherosclerotic vascular disease. Consistent with this, irradiated mice transplanted with bone marrow cells from *Hmox1*^*-/-*^ mice show enhanced atherosclerotic lesions together with a greater macrophage content and increased inflammatory cytokines including monocyte chemotactic protein 1 [16].

The whole genome duplication event in the teleost lineage resulted in two homologs (*hmox1a* and *hmox1b*) of human *HMOX1* in zebrafish genome. Both genes are inducible under oxidative stress conditions. However, *hmox1a* has been shown to be expressed at a significantly higher level during the early developmental stage and in response to oxidative stress conditions than *hmox1b* [17, 18].

Until now, no *hmox1a* mutant zebrafish allele has been generated to study the function of heme oxygenase in zebrafish. Using a combination of CRISPR-Cas9 and TALEN mutagenesis techniques, we have studied the function of zebrafish *hmox1a* in development and leukocyte biology.

## Methods

### Zebrafish husbandry

Adult zebrafish were maintained at Centenary Institute and embryos were obtained by natural spawning followed and raised in E3 media at 28-32°C (Sydney Local Health District AWC Approvals: 16-037 and 17-036). All work with the mutant allele and the *Tg(mpeg1:gfp)* transgenic line were performed in a WT TU/AB genetic background, CRISPR work with the *Tg(mfap4:turquoise)* transgenic line was performed in a WT AB genetic background.

### Generation of a *hmox1a* mutant allele by TALEN mutagenesis

The CHOPCHOP web tool was used to predict TALEN pairs targeting *hmox1a*. The selected TALEN information is listed in Supplementary Table 1. Constructs were assembled using Golden Gate assembly and cloned into pCS2TAL3DDD and pCS2TAL3RRR vectors [19-21].

Genome editing was performed as described [21]. Briefly, TALENs, used to generate mutant zebrafish, were constructed using the Golden Gate assembly method [19, 20]. mRNAs were synthesized using the Transcription Kits (Invitrogen) and diluted to 50 ng/μL for microinjection. Samples were prepared immediately before each injection. mRNAs were injected into zebrafish embryos at the early one-cell stage to 10% of the volume of the cell (∼1 nL). Injected embryos were raised until they reached adulthood and then outcrossed to wild-type fish. Embryos from these crosses were used for screening to identify mutants. The resulting mutant line is *hmox1a*^*vcc42*^.

After initial genotyping characterisation, experimental work was carried out on embryos derived from F3-F5+ generation adults.

### Genotyping

For *hmox1a*^*vcc42*^ genotyping, a piece of fin was cut off from larval or adult zebrafish for DNA extraction. PCR was performed with the *hmox1a* genotyping primers described in Supplementary Table 2 using an annealing temperature of 60°C and an extension time of 30 seconds. PCR products were digested by *HinfI* restriction enzyme at 37°C for 8-24 h and the digests were resolved by agarose gel electrophoresis.

### Phenotyping

Embryos showing no eyes, severe cyclopisa, small eyes, small trunk somites and heart, reduced notochord or missing floor plate were described as ‘abnormal’. Embryos showing no gross abnormalities were described as ‘normal’.

### CRISPR/Cas9 Gene Editing Technique

The gRNA target sites for each gene were designed using https://www.crisprscan.org, gRNA oligonucleotide sequences are listed in Supplementary Table 3. The gRNA templates were amplified by PCR with scaffold reverse primer then were transcribed with HiScribe™ T7 High Yield RNA Synthesis Kit (NEB) [22]. One cell stage embryos were injected with 1 nL of a pre-incubated mixture containing 200 ng/μL of the four gRNAs, or eight gRNAs in the double knockdown, and 2 ng/μL Cas9.

### Imaging

Imaging was carried out on Leica M205FA and DM6000B, and Deltavision Elite microscopes as previously described [23-25].

### Drug treatments

Copper-protoporphyrin IX (CuPP) (Frontier) and tin-protoporthyrin IX (SnPP) (Frontier) were dissolved in 0.05M NaOH and then diluted in pH 8.0 Tris-HCl solution to a stock concentration of 10 mM. All compounds were used at a final concentration of 10 μM. Embryos were treated with drugs within 3 h of egg fertilization and changed every three days for developmental studies or immediately after tail wounding and not changed for the duration of the macrophage migration assays.

### Quantitative Real–time PCR

Total RNA was extracted from homogenates using Trizol (Thermofisher) and cDNA was synthesized with the High Capacity cDNA Synthesis Kit (Applied Biosystems). qPCR reactions were performed on a LightCycler^®^ 480 System. Gene expression was quantified by the delta–delta C_T_ method normalized to host *bact*. Sequences of primers are listed in Supplementary Table 2.

### Tail wounding assay

Caudal fin amputation was performed on 4 dpf embryos. Zebrafish larvae were anaesthetized using tricaine for wounding and imaging experiments. Embryos were cut posterior to the notochord and then recovered to fresh E3 and kept at 28°C. Macrophages were imaged at 6 and 24 h post wounding (hpw) after tail-fin incision of fluorescent reporter zebrafish larvae.

### Statistics

All ANOVA and Student’s *t* tests as appropriate for the number of comparisons were carried out using Graphpad Prism. Each data point indicates a single animal unless otherwise stated.

## Results

### Generation of zebrafish *hmox1a*^*vcc42*^ mutant allele

The traditional method to establish a mutant animal model is to induce a premature stop codon caused by nucleotide deletion. In several cases however, phenotypes in knockout mutant animals using this method have been shown to lack specificity compared to antisense knockdown methods [26-29]. Recently, it was determined that nonsense-mediated, decay-induced genetic compensation can cause different phenotypes between knockout mutants and knockdown methods [26]. Animal models established with in-frame mutation or mutant targeting at the gene promoter area have a reduced probability of genetic compensation [26]. To minimize the potential effects of genetic compensation by the paralogs *hmox1b, hmox2a*, and *hmox2b*, we generated an in-frame mutation allele of *hmox1a* targeting the ATG start codon in exon 2 (Fig. 1A).

**Figure 1.**
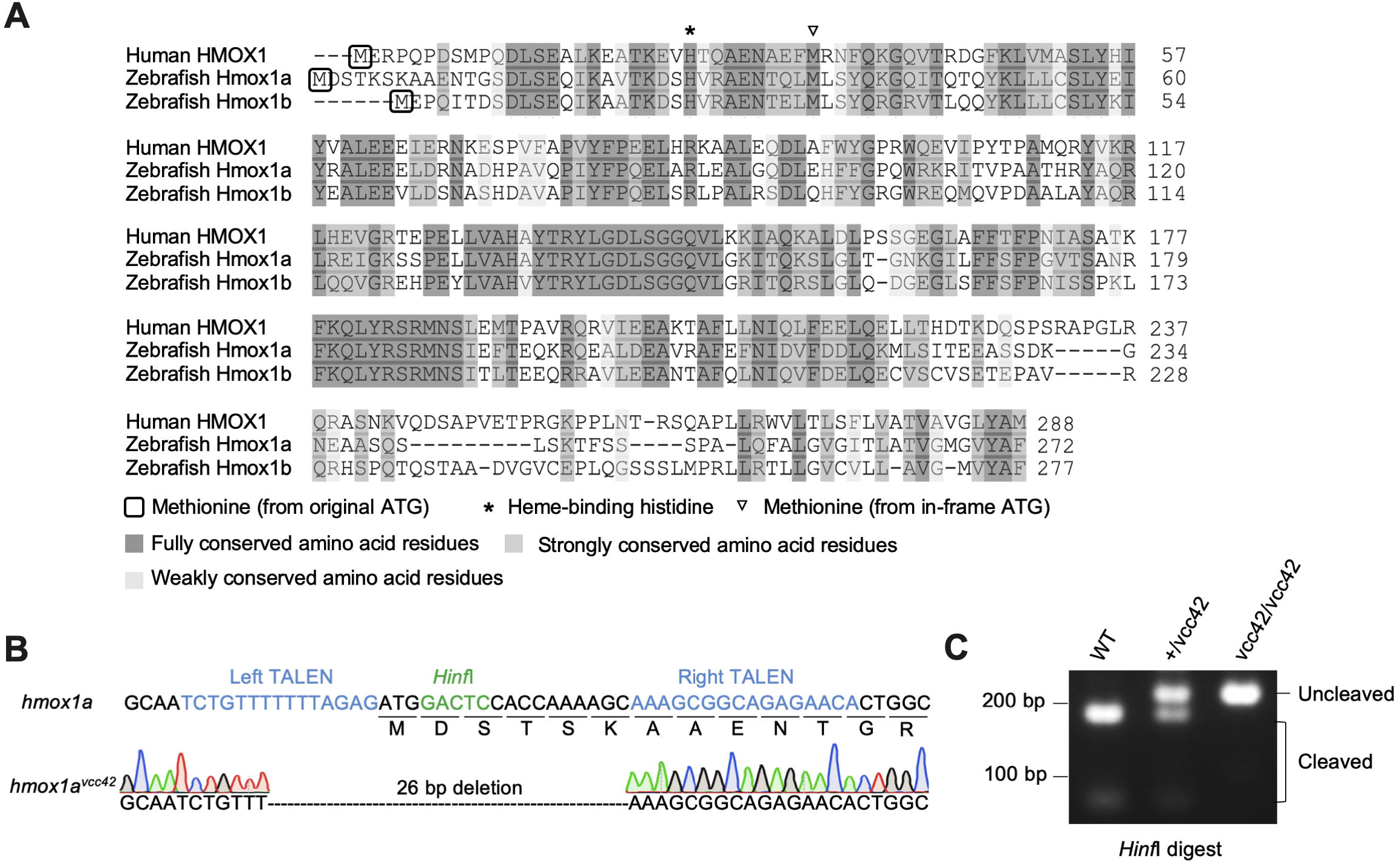
Creation of the *hmox1a*^*vcc42*^ allele. **A**. Alignment of *HMOX1* homologs. Dark grey background indicates amino acid residues that are identified in all three proteins (Uniprot). Light grey background indicates amino acid residues that are similar in all four proteins (Uniprot). The purple background indicates the functional histidine. Similarity, identity and functional histidine were predicted by Uniprot database (https://www.uniprot.org/align/). **B**. Partial sequence of the *hmox1a*^*vcc42*^ allele. TALEN mRNA recognition sequences are blue, and the *HinfI* site in the spacer region is highlighted in green. Exons are indicated by grey-filled boxes with numbers. Black trace indicates guanine, blue trace indicates cytosine, green trace indicates adenine and red trace indicates thymine. **C**. Representative results from PCR-based genotyping using DNA extracted from the fins of *hmox1a*^*vcc42*^ adult zebrafish followed by *HinfI* digests.

An in-frame ATG was found in exon 3 at amino acid 37, which was predicted to be active when the initial ATG was deleted. The translated amino acid fragment from the second ATG start site in the mutant mRNA sequence was predicted to be non-functional as the heme-binding histidine is absent in the predicted amino acid sequence (Fig. 1A).

To generate the mutant allele, TALEN-injected embryos were raised to adulthood and three F0 fish with mutations in somatic tissue were individually outcrossed with wild-type (WT) fish. After sequencing of F1 fish, we identified an in-frame 26 bp deletion mutant *hmox1a* allele with deletion of the original start codon, resulting in a predicted mutant protein lacking the functional histidine (Fig. 1B). The mutant allele was designated as *hmox1a*^*vcc42*^, and offspring were propagated for analysis (Fig. 1C).

### Characterization of *hmox1a*^*vcc42*^ expression

Genomic sequencing revealed the ‘AG’ splice acceptor site preceding exon 2 was deleted in *hmox1a*^*vcc42*^. Because there was no splice acceptor attached at the beginning of exon 2 to define the termination of an intron in front of it, the mutant exon 2 would be treated as a part of a long intron till the next splice acceptor before exon 3 (Fig. 2A). Thus, we hypothesized that mature mutant *hmox1a*^*vcc42*^mRNA would skip exon 2.

**Figure 2.**
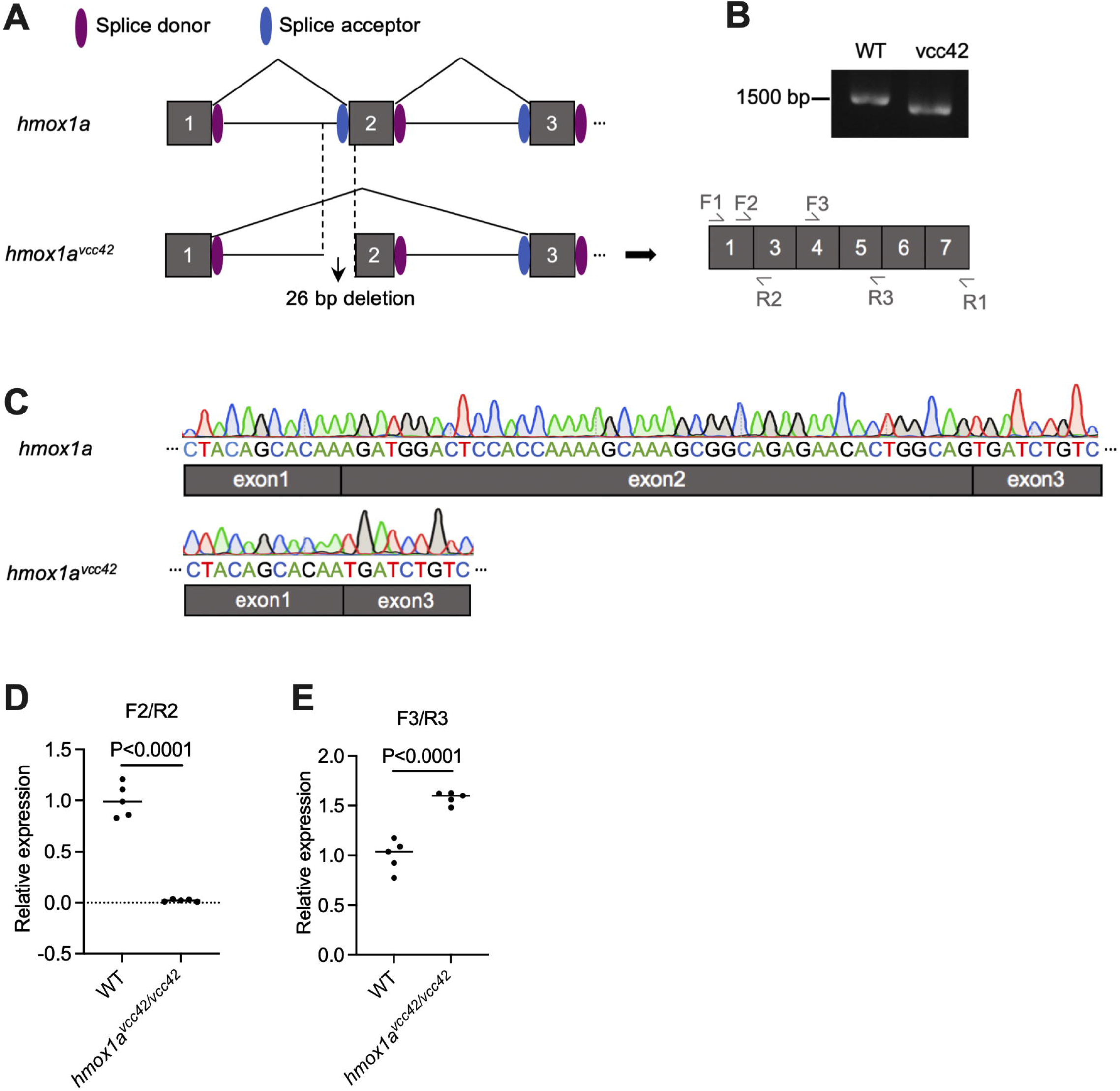
Characterization of the *hmox1a*^*vcc42*^ allele mRNA transcript. **A**. Schematic diagram of the (predicted) splicing process. Primers used for whole-transcript PCR amplification are indicated as F1 and R1. Primers F2/R2 and F3/R3 were used for qPCR analysis. **B**. DNA gel electrophoresis visualization of the exon 1-7 amplicon of mutant *hmox1a* mRNA. **C**. Alignment of the mRNA transcript of the mutant *hmox1a*^*vcc42*^ allele sequence (bottom) with WT zebrafish *hmox1a* sequence (top). **D**. Quantification of *hmox1a* exon 2 mRNA by RT-qPCR analysis. **E**. Quantification of *hmox1a*^*vcc42*^ exons 4-5 mRNA by RT-qPCR analysis. Statistical testing by Student’s *t* test, error bars represent one standard deviation, and each data point represents a biological replicate of pooled embryos.

To explore this hypothesis, RNA was isolated from homozygous *hmox1a*^*vcc42/vcc42*^ and WT zebrafish and cDNA was synthesized via reverse transcription. Primer F1 targeting the start of exon 1 was used as forward primer and primer R1 targeting the end of exon 7 was used as reverse primer (Fig. 2A). The mutant *hmox1a* exon 1-7 amplicon was slightly smaller than WT as expected from the 41 bp size of exon 2 (Fig. 2B). The exon skipping in *hmox1a* mRNA was further confirmed by sequencing the cDNA amplicon (Fig. 2C).

To confirm the effect of the *hmox1a*^*vcc42*^ allele on *hmox1a* transcripts, qPCR analysis was performed to examine the expression of sections of the *hmox1a* mRNA. Primers spanning the exon 1-2 and 2-3 junctions were chosen for the first analysis (F2/R2; Fig. 2A). As expected, *hmox1a* exon 2 junction-containing mRNA was almost undetectable in the mutants (Fig. 2D).

Conversely, it was hypothesized that other exon sequences, including exon 1 and exon 3-7, should still maintained in *hmox1a*^*vcc42*^ transcripts. To investigate this hypothesis, primers (F3/R3; Fig. 2A) targeting exon 4 and exon 5 regions were used for qPCR analysis at the same time. Mutant exons 4 and 5 of *hmox1a* mRNA were sustained and not degraded (Fig. 2E), indicating that nonsense-mediated decay is not triggered by the *hmox1a*^*vcc42*^ allele.

### Morphologic analysis of *hmox1a*^*vcc42/vcc42*^ zebrafish

The percentage of homozygous *hmox1a*^*vcc42/vcc42*^ embryos generated from *hmox1a*^*vcc42*^ heterozygote in crosses slightly decreased from 3 to 7 dpf and further gradually decreased to 30 dpf; from 30 dpf onwards, the percentage of homozygous mutant embryos was stable (Table 1).

To investigate the reason for the loss of homozygous mutants, 3 dpf embryos from sibling matched *hmox1a*^*WT/WT*^ and *hmox1a*^*vcc42/vcc42*^ adult in crosses were collected for morphologic analysis. Embryos showing no eye, severe cyclopia or small eyes, trunk somites and heart, reduced notochord and missing floor plate were described as ‘abnormal’ (Fig. 3A). Embryos that showed no gross abnormalities were described as ‘normal’. While WT embryos did not display abnormal morphology (0%, n= 368), around 20% of *hmox1a*^*vcc42/vcc42*^ embryos displayed an abnormal phenotype with shorter length at 3 dpf (Fig. 3B). The percentage of dead *hmox1a*^*vcc42/vcc42*^ embryos at 11 dpf correlated with the proportion of abnormal *hmox1a*^*vcc42/vcc42*^ embryos at 3 dpf (Fig. 3C, Table 2). This analysis suggests that a minority of zebrafish *hmox1a*^*vcc42/vcc42*^ embryos undergo an aberrant morphogenetic process and are unable to survive embryogenesis.

**Figure 3.**
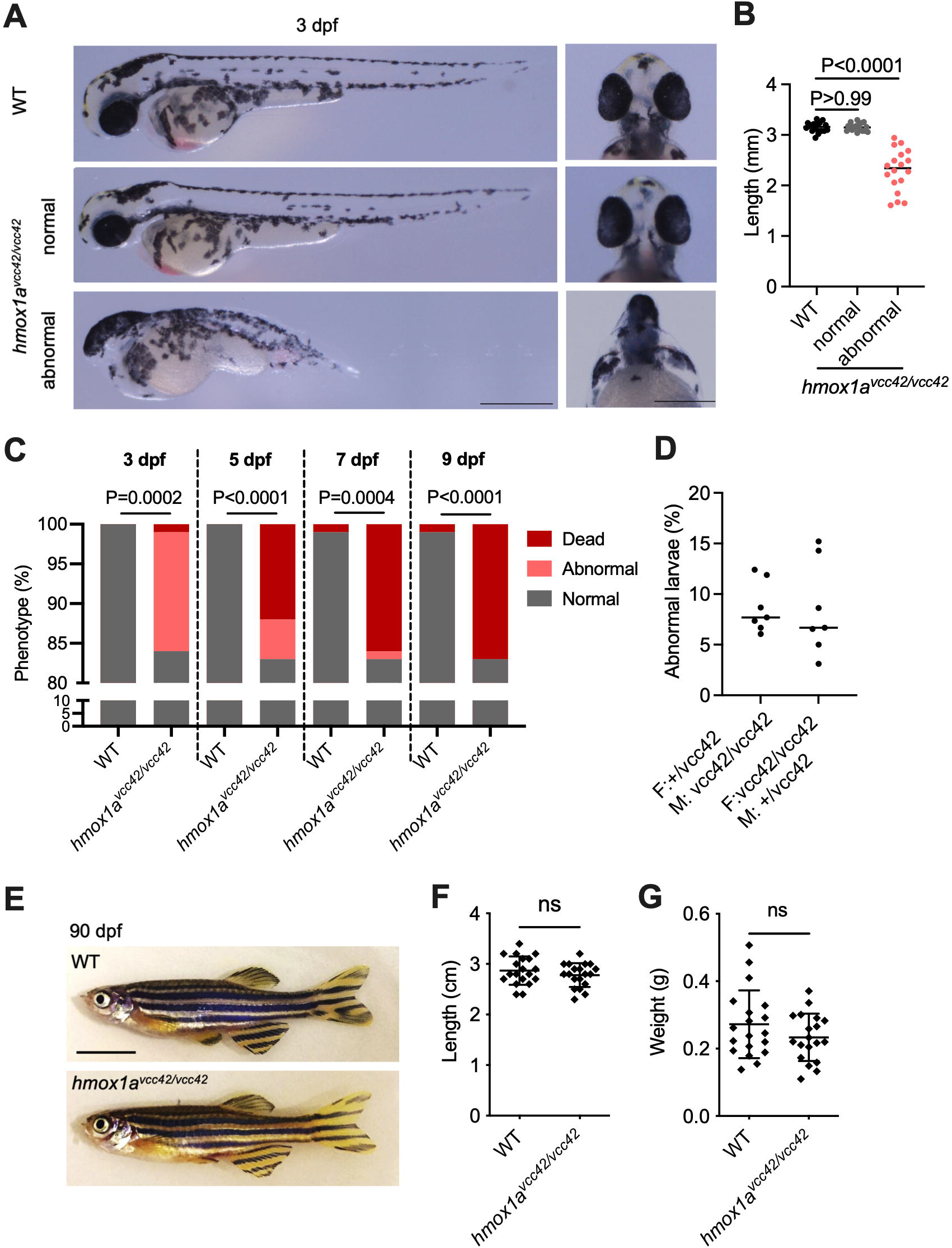
Morphologic analysis of *hmox1a*^*vcc42/vcc42*^ embryos. **A**. Images of 3 dpf embryos from a *hmox1a*^*+/vcc42*^ intercross. Abnormal embryos displayed apparent shorter lengths. Scale bar represents 0.5 mm. **B**. Quantification of 3 dpf embryo length in WT and *hmox1a*^*vcc42/vcc42*^ normal and abnormal embryos. Statistical testing carried out by ANOVA. Data is pooled from 3 WT and 4 *hmox1a*^*vcc42/vcc42*^ replicates. **C**. Proportions of abnormal embryo morphology and death during development. Raw counts are available in Table 2. Statistical testing was performed using Chi-squared test on raw counts with abnormal and dead embryos treated as one group. **D**. Proportions of abnormal embryos scored at 3 dpf from female *hmox1a*^*+/vcc42*^; male *hmox1a*^*vcc42/vcc42*^ (F: +/vcc42; M: vcc42/vcc42) and female *hmox1a*^*vcc42/vcc42*^; male *hmox1a*^+/vcc42^ (F: vcc42/vcc42; M: +/vcc42) crosses. Statistical testing was performed using *t* test, and each data point represents a biological replicate of pooled embryos from an individual crossing. **E**. Representative images of adult (90 dpf) WT and *hmox1a*^*vcc42/vcc42*^ zebrafish from an *hmox1a*^*+/vcc42*^ intercross. Scale bar represents 0.5 cm. **F**. Quantification of the length (left) and weight (right) of adult (90 dpf) *hmox1a*^*vcc42*^ zebrafish from an *hmox1a*^*+/vcc42*^ intercross. Statistical testing was performed using *t* test, and error bars represent SEM.

It is known that embryo morphogenetic development is not only affected by gene expression but also depends on maternal factors that are required for processes prior to the activation of the zygotic genome. To identify whether *hmox1a* is a maternal factor, embryos from *hmox1a*^*vcc42/vcc42*^ females crossed with *hmox1a*^*+/vcc42*^ males and *hmox1a*^*+/vcc42*^ females crossed with *hmox1a*^*vcc42/vcc42*^ males were screened separately. No differences were observed between these crosses (Fig. 3D), suggesting that the morphological phenotype was independent of the paternal genotype.

The 20% of *Hmox1*-deficient mice that survive until adulthood display smaller size and less body weight [30]. To investigate whether zebrafish *hmox1a* deficiency also affects adult morphology, *hmox1a*^*vcc42/vcc42*^ adult zebrafish were assessed for morphology, body length and weight. In contrast to *Hmox1*^*-/-*^ mice, *hmox1a*^*vcc42/vcc42*^ adult zebrafish developed without gross abnormalities compared to their WT clutch mates (Fig. 3E). Body length and weight was also comparable between *hmox1a*^*vcc42/vcc42*^ and WT clutch mate adults (Fig. 3F). These results indicate zebrafish *hmox1a* may have a different impact on embryogenesis and development compared to mouse *Hmox1*, or that genetic compensation by another Hmox family member plays an important compensatory role during the development of *hmox1a*^*vcc42/vcc42*^ zebrafish.

### Zebrafish *hmox1b* partially compensates for loss of *hmox1a* during embryonic development

As a consequence of the whole genome duplication event in zebrafish, there are additional paralogs *hmox1b, hmox2a*, and *hmox2b* that may increase the survival of *hmox1a*^*vcc42/vcc42*^ zebrafish compared to mice, where *Hmox2* is the only related gene to *Hmox1*.

We first examined the expression of *hmox1b, hmox2a* and *hmox2b* in *hmox1a*^*vcc42/vcc42*^ zebrafish. Significant upregulation of *hmox1b* was detected in phenotypically normal *hmox1a*^*vcc42/vcc42*^ embryos compared to WT embryos and *hmox1b* expression was even higher in abnormal *hmox1a*^*vcc42/vcc42*^ embryos than the other phenotypically normal embryos of either genotype (Fig. 4A). The zebrafish *hmox2* genes, *hmox2a* and *hmox2b*, were expressed at a slightly higher level in abnormal *hmox1a*^*vcc42/vcc42*^ embryos relative to WT embryos, and there was no change in the expression of either *hmox2a* or *hmox2b* in normal homozygous *hmox1a*^*vcc42/vcc42*^ embryos (Fig. 4A). Upregulation of *hmox1b*, but not of *hmox2a* or *hmox2b*, was also seen in *hmox1a* crispants (Fig. 4B).

**Figure 4.**
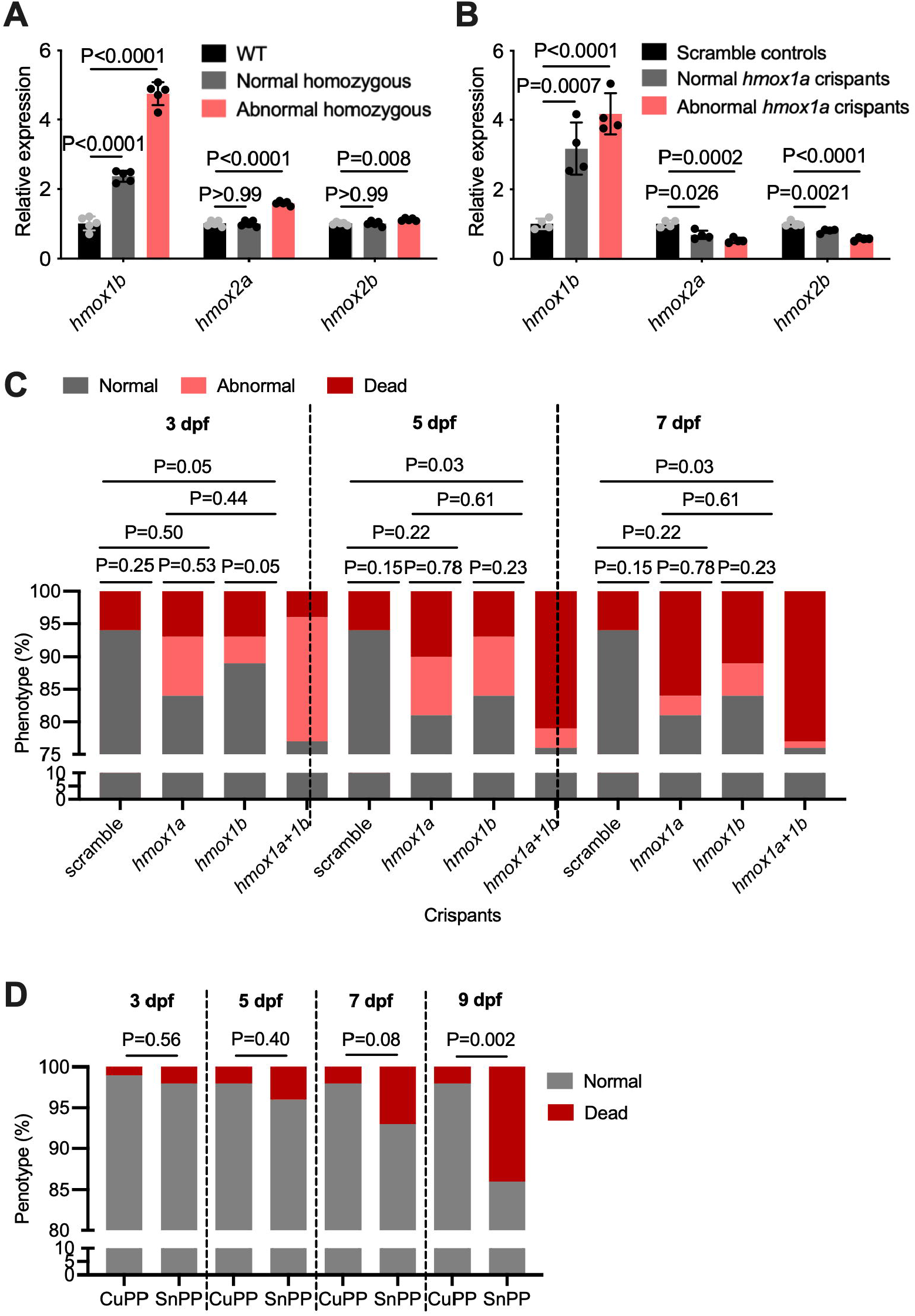
*hmox1a* and *hmox1b* have non-redundant roles in zebrafish development. **A**. Quantification of *hmox1b, hmox2a* and *hmox2b* expression in 3 dpf WT and *hmox1a*^*vcc42/vcc42*^ normal and abnormal embryos. Statistical testing was performed using the one-way ANOVA, error bars represent one standard deviation, and each data point represents a biological replicate of at least 10 pooled embryos. **B**. Quantification of *hmox1b, hmox2a* and *hmox2b* expression in 3 dpf scramble control and *hmox1a* crispant normal and abnormal embryos. Statistical testing was performed using the one-way ANOVA, error bars represent one standard deviation, and each data point represents a biological replicate of at least 10 pooled embryos. **C**. Quantification of abnormal embryo morphology and death during double *hmox1a* and *hmox1b* crispant development. Statistical testing was performed using Chi-squared test on raw counts with abnormal and dead embryos treated as one group. **D**. Quantification of abnormal embryo morphology and death during development in embryos treated with SnPP. Raw counts for panels C and D are available in Table 2.

Knockdown of *hmox1a* caused the appearance of visually similar developmental defects to those seen in homozygous *hmox1a*^*vcc42/vcc42*^ embryos however there was a basal rate of pre-3 dpf embryo death caused by the microinjection that obscured any statistically significant difference in rate of developmental defects in *hmox1a* crispants (Fig. 4C, Table 2). Knockdown of *hmox1b* also caused some developmental defects similar to the loss of *hmox1a* but at a statistically similar rate to the embryo death in scrambled controls and overall developmental defects in *hmox1a* crispants (Fig. 4C, Table 2). Double knockdown of *hmox1a* and *hmox1b* by CRISPR-Cas9 increased the rate of developmental defects compared to scramble control embryos across 3-7 dpf and increased the rate of developmental defects compared to *hmox1b* knockdown embryos at 3 dpf (Fig. 4C, Table 2).

Treatment of developing WT embryos with the Hmox inhibitor tin protoporphyrin (SnPP) caused increased mortality after 9 dpf compared to copper protoporphyrin (CuPP) controls (Fig. 4D, Table 2). Together, these results demonstrate a conserved role for zebrafish Hmox1a and Hmox1b in embryonic development.

### Conservation of Hmox1a regulation of macrophage migration

To investigate the role of *hmox1a* in macrophage biology, we crossed the *hmox1a*^*vcc42*^ allele into the *Tg(mpeg1:GFP)*^*vcc7*^ background.

As *Hmox1* has been reported to be essential for hematopoiesis [31, 32], the total fluorescent area of macrophages was measured by fluorescence microscopy in 4 dpf *Tg(mpeg1:GFP)*^*vcc7*^ zebrafish embryos, where macrophages are marked by GFP. No difference was observed between the WT control and *hmox1a*^*vcc42/vcc42*^ larvae (Fig. 5A).

**Figure 5.**
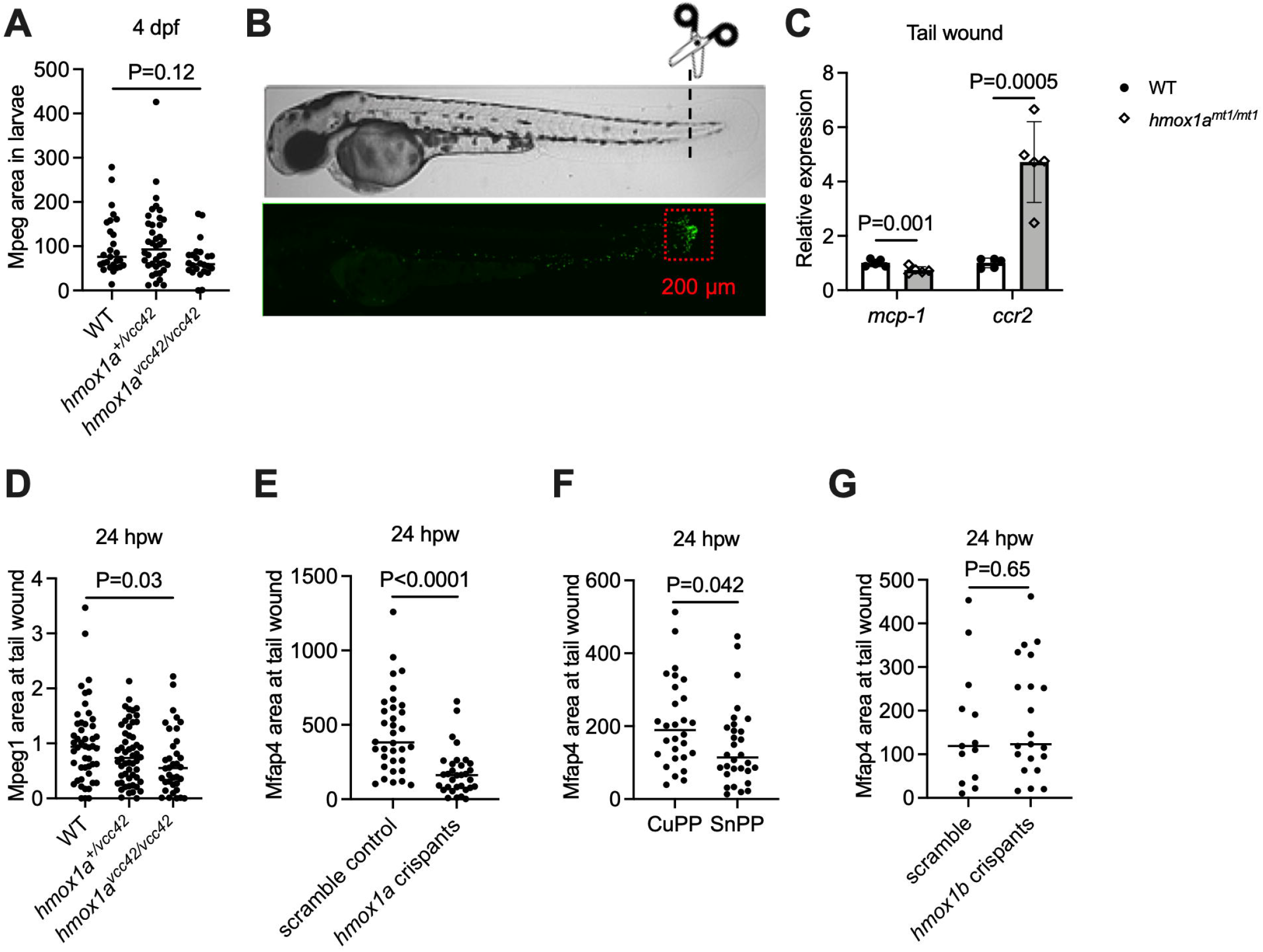
Zebrafish *hmox1a* aids macrophage migration. **A**. Quantification of total macrophages of *hmox1a*^*vcc42/vcc42*^, *hmox1a*^*+/vcc42*^ and WT larvae. Statistical testing was performed by one-way ANOVA. **B**. Images represent embryos at 4 dpf (top) and macrophage recruitment to wound site (bottom) at 6 hpw, red box indicates area measured for macrophage recruitment in panels D-G. **C**. Relative quantity of *mcp-1* and *ccr2* transcripts measured by qPCR at 6 hpw, and each datapoint represents biological replicate of a pool of at least 10 embryos. **D**. Quantification of macrophage accumulation at wound site by normalized fluorescent area. Data are pooled from two biological replicates. Statistical testing was performed by one-way ANOVA. **E**. Quantification of macrophages at wound site by fluorescent area in scramble control larvae and *hmox1a* crispants. **F**. Quantification of macrophages at wound site by fluorescent area in PPIX or SnPP treated zebrafish larvae. **G**. Quantification of macrophages at wound site by fluorescent area in scramble control larvae and *hmox1b* crispants. Statistical testing was performed by *t* test unless otherwise stated, error bars represent one standard deviation, and data are from one experiment representative of at least two biological replicates unless otherwise stated.

Chemokine production plays a crucial role in the recruitment of leukocytes to sites of inflammation. The monocyte chemoattractant protein-1 (MCP-1) and its receptor CCR2 are regulated by HMOX1 in mouse models of mycobacterial infection [33, 34]. To determine if the relationship between Hmox1 and macrophage chemoattractant production is conserved in zebrafish, we performed tail wounding of zebrafish embryos (Fig. 5B). The expression of the chemokine *mcp-1* and receptor *ccr2* were examined at 6 h post wounding (hpw) by qPCR. Expression of *mcp-1* was reduced while *ccr2* expression was increased in *hmox1a*^*vcc42/vcc42*^ larvae compared to WT littermates (Fig. 5B).

To investigate the functional consequence of reduced *mcp-1* expression, we quantified macrophage migration to the wound. There was decreased macrophage accumulation at wound area in *hmox1a*^*vcc42/vcc42*^ homozygous larvae compared to their WT littermates at 24 hours post wounding (Fig. 5D).

CRISPR/Cas9 technology was used to knockdown *hmox1a* expression in *Tg(mfap4:turquiose)*^*xt12*^ zebrafish embryos, where the *mfap4* promoter drives macrophage-specific expression comparable to the *mpeg1* promoter [35]. The number of wound-associated *mfap4*:Turquiose^+^ cells was significantly reduced in *hmox1a* crispants compared to scramble control larvae (Fig. 5E).

To confirm these genetic results, the HMOX inhibitor, SnPP, was used to treat tail wounded WT *Tg(mfap4:turquiose)* ^*xt12*^ larvae. SnPP decreased macrophage number at the wound site compared to CuPP-treated controls (Fig. 5F).

As *hmox1b* knockdown had produced some of the developmental defects seen in *hmox1a* depletion, we examined the effect of *hmox1b* knockdown on macrophage migration to a wound. Knockdown of *hmox1b* did not affect macrophage migration (Fig. 5G).

These results demonstrate depletion of *hmox1a*, but not *hmox1b*, decreases macrophage migration in zebrafish.

## Discussion

We report the generation and characterization of a zebrafish with a *hmox1a* mutant allele with approximately 20% lethality in homozygotes during embryonic development. Morphologic analysis of *hmox1a*^*vcc42/vcc42*^ embryos revealed the occurrence of developmental disorders from 3 dpf that progressed to lethality within 14 dpf. These developmental defects were reproduced in double *hmox1a* and *hmox1b* crispants, and mortality was phenocopied by Hmox1 inhibitor treatment. Using live imaging and gene expression analyses we demonstrate conservation of the link between *hmox1a* and macrophage migration in zebrafish. This *hmox1a*^*vcc42*^ allele provides a platform to investigate the role of *hmox1a* in response to different environmental stresses in zebrafish.

The first *Hmox1*^-/-^ mouse model was established by Poss *et al* in 1997 with total deletion of exons 3 and 4, and partial deletion of exon 5. About 80% of *Hmox1*^-/-^ mice succumbed to prenatal lethality and premature mortality [30]. Our *hmox1a*^*vcc42*^ allele, which only contains deletion of exon 2, caused disorders of development and premature mortality in about 20% of *hmox1a*^*vcc42/vcc42*^ embryos. This ∼20% mortality rate was similar in *hmox1a* crispants suggesting that the different survival rates between *Hmox1*^-/-^ mice and *hmox1a*-depleted zebrafish is not caused by residual peptide translated from the *hmox1a*^*vcc42*^ transcript. A possible explanation for the difference in survival is the high expression of compensatory *hmox1b*, which exists in zebrafish but not in mice. Our double knockdown experiment where depletion of *hmox1a* and *hmox1b* by CRISPR-Cas9 mutagenesis increased the rate of developmental abnormalities compared to control and *hmox1b*-depleted, but not *hmox1a*-depleted, embryos suggest *hmox1a* has a more important role in zebrafish development than *hmox1b*.

The upregulation of *hmox1b* in *hmox1a*^*vcc42/vcc42*^ embryos might be caused by genetic compensation resulting from the genomic editing in the *hmox1a* gene locus. Although genetic compensation has been described for a long time [36, 37], the nonsense-mediated decay pathway mechanism has only been recently described [26, 38]. Mutant alleles that avoid induction of the premature stop codon, such as the *hmox1a*^*vcc42*^ allele, have been reported to reduce the probability of inducing genetic compensation [39, 40]. We also observed upregulation of *hmox1b* in *hmox1a* crispants. This suggests the upregulation of *hmox1b* in *hmox1a*^*vcc42/vcc42*^ embryos is more likely to be caused by the stress within embryos secondary to the loss of Hmox1a function. Although we observed some developmental defects in *hmox1b* crispants, the lack of statistical significance compared to scramble control embryos and lack of significant additional developmental defects in double knockdown embryos compared to *hmox1a* knockdown alone suggests a biologically insignificant function of *hmox1b* in zebrafish embryogenesis.

About 80% of *hmox1a*^*vcc42/vcc42*^ larvae survived until adulthood without gross abnormalities at 90 dpf. This is in accordance with the mouse study where, in the minority of surviving *Hmox1*-deficient mice, no difference in body weight was reported between *Hmox1*-deficient homozygous and heterozygous mice until the age of 20 weeks when body size started to be lost in *Hmox1*-deficient mice [30]. Further morphological analysis of *hmox1a*^*vcc42/vcc42*^ zebrafish over 90 dpf is necessary to investigate the conservation of this effect in zebrafish.

It has been reported that expression of MCP-1, also known as CCL2, was significantly increased in the liver, spleen, bronchoalveolar lavage fluid (BALF) and serum samples of *M. avium* infected *Hmox1*^*-/-*^ mice compared to WT controls [34, 41]. However, inhibition of HMOX1 in *M. avium*-infected RAW 264.7 cells reduced the expression of *MCP-1* compared to untreated cells [34]. The different results from reported studies may be due to the examination of MCP-1 expression at different levels since in the two experiments using *Hmox1*^*-/-*^ mice, MCP-1 expression was analyzed at protein level [34, 41], but in the macrophage infection experiment, the gene expression of *MCP-1* was analyzed [34]. In our study, decreased *mcp-1* transcript levels were detected in tail wounded *hmox1a*^*vcc42/vcc42*^ larvae, which was consistent with the *in vitro* macrophage study [34]. Taken together, these results suggest that *mcp-1* expression is dependent on *hmox1a* expression in zebrafish and raise the possibility that macrophage function is altered by increased alternative activation of macrophages driven by the Ccl2/Ccr2 axis.

Using live imaging in zebrafish embryos, we provide direct evidence linking Hmox1a to the regulation of macrophage migration. We found a macrophage migration defect in *hmox1a*^*vcc42/vcc42*^ larvae that was recapitulated in *hmox1a* crispants and SnPP-treated larvae, but not in *hmox1b* crispants. This was similar to the smaller effect of *hmox1b* depletion on development and provides further evidence that *hmox1a* provides the most conserved functions expected of heme oxygenase 1 in zebrafish embryos. Further work with loss of function alleles of the zebrafish *hmox* family are necessary to determine which gene or genes mediate the expected functions of heme oxygenase 1 in other biological processes.

## Acknowledgements

We thank Drs Elinor Hortle and Pradeep Cholan for zebrafish infection and analysis methods training, Drs Angela Kurz and Kristina Jahn of Sydney Cytometry for assistance with imaging equipment, the Victor Chang Cardiac Research Institute BioCORE staff for maintenance of zebrafish, and members of the Tuberculosis Research Program at the Centenary Institute for helpful discussion.

