## Supplementary Tables for "Zebrafish heme oxygenase 1a is necessary for normal development and macrophage migration"

Supplementary Table 1: Information of chosen TALEN assembles

| Target sequence | Genomic location | TALE 1 | TALE 2 | Off-targets | Restriction sites |
| --- | --- | --- | --- | --- | --- |
| TGTTCTCTGCCGCTTT*GCTTTTGGTGGAGTCCA*TCTCTAAAAAAACAGA | chr3:25886359 | NG NN NG NG HD NG HD NG NN HD HD NN HD NG NG NG | NG HD NG NN NG NG NG NG NG NG NG NI NN NI NN NI | 0 | *Hinf*I |

Supplementary Table 2: Sequences of PCR primers

| Gene target and purpose | Forward primer | Reverse Primer |
| --- | --- | --- |
| *hmox1a* qPCR F1/R1 | 5’– ACCCTCTCTGCTTTGTCATGAGAAA | 5’– CGGTTGGCATGGGAGTTTACGCTTTT |
| *hmox1a* qPCR F2/R2 | 5’–GGGCAGGACTTGGAGCACTT | 5’–GGACTGCTCTTGCCAATCTCT |
| *hmox1a* qPCR F3/R2 | 5’–TAAAAACGAAGTGGGGCGGT | 5’–TGTTCAGACAGATCACTGCCA |
| *hmox1a* genotyping | 5’-AATCTGAGTCTGCGGAAATGGCGACATTTGATAACAAATAGGGCAATCTGT | 5’-AATCTGAGTCTGCGGAAATGGCGACATTTGATAACAAATAGGGCAATCTGT |
| *hmox1b* qPCR | 5’–AGCTACCAGAGGGGCCGAGT | 5’–CGCCTCGTAGATCTTGTAGAGC |
| *hmox2a* qPCR | 5’–ATGGCGGTCAGTGGAAACACAACC | 5’–GGCAACAGCAGCAACCAATGTGGC |
| *hmox2b* qPCR | 5’–TTTAGGAGGTTGAGTTGGAGTCAG | 5’–TTCTGCCTTCTGGTGCACTTCT |
| *ccr2* qPCR | 5’–GCAACAATGGCAACGCAAAG | 5’–GTGAGCCCAGAACGGAAGTG |
| *mcp-1* qPCR | 5’–GTCTGGTGCTCTTCGCTTTC | 5’–TGCAGAGAAGATGCGTCGTA |

Supplementary Table 3: Primer sequences used for CRISPR-Cas9 gRNA generation

| Primer name | Sequence |
| --- | --- |
| *hmox1a* target 1 | TAATACGACTCACTATAGGACTCCACCAAAAGCAAAGGTTTTAGAGCTAGAA |
| *hmox1a* target 2 | TAATACGACTCACTATAGGGTGTTTTCAGCTCTGACGGTTTTAGAGCTAGAA |
| *hmox1a* target 3 | TAATACGACTCACTATAGGAGATCTACCGAGCGCTGGGTTTTAGAGCTAGAA |
| *hmox1a* target 4 | TAATACGACTCACTATAGGTAAATGGGCTGCACTGCTGTTTTAGAGCTAGAA |
| *hmox1b* target 1 | TAATACGACTCACTATAGGGTCACTGTCGGTGATCTGGTTTTAGAGCTAGAA |
| *hmox1b* target 2 | TAATACGACTCACTATAGGAGCGTCACTCGGCCCCTCGTTTTAGAGCTAGAA |
| *hmox1b* target 3 | TAATACGACTCACTATAGGTCTACGAGGCGCTGGAGGGTTTTAGAGCTAGAA |
| *hmox1b* target 4 | TAATACGACTCACTATAGGTGGGAGCCACGGCATCATGTTTTAGAGCTAGAA |
| scramble target 1 | TAATACGACTCACTATAGGCAGGCAAAGAATCCCTGCCGTTTTAGAGCTAGAAATAGC |
| scramble target 2 | TAATACGACTCACTATAGGTACAGTGGACCTCGGTGTCGTTTTAGAGCTAGAAATAGC |
| scramble target 3 | TAATACGACTCACTATAGGCTTCATACAATAGACGATGGTTTTAGAGCTAGAAATAGC |
| scramble target 4 | TAATACGACTCACTATAGGTCGTTTTGCAGTAGGATCGGTTTAGAGCTAGAAATAGC |
| Scaffold | AAA AGC ACC GAC TCG GTG CCA CTT TTT CAA GTT GAT AAC GGA CTA GCC TTA TTT TAA CTT GCT ATT TCT AGC TCT AAA AC |
