## Supplementary material for "Zebrafish heme oxygenase 1a is necessary for normal development and macrophage migration": Tables

Table 1. Mendelian distribution of *hmox1a^vcc42^* in embryos from *hmox1a*^+/vcc42^ in crosses.

| Date | WT | +/vcc42 | Vcc42/vcc42 | Total n |
| --- | --- | --- | --- | --- |
| Day 3 | 17 | 29 | 14 | 60 |
| Day 7 | 19 | 29 | 12 | 60 |
| Day 30 | 25 | 57 | 18 | 100 |
| Day 90 | 25 | 49 | 16 | 90 |

Table 2: Phenotyping of embryos during development

|  | Number of normal embryos; abnormal embryos; dead embryos | | | | | | | | | | |
| --- | --- | --- | --- | --- | --- | --- | --- | --- | --- | --- | --- |
| Experimental condition | 1dpf | 2dpf | 3dpf | 4dpf | 5dpf | 6dpf | 7dpf | 8dpf | 9dpf | 10dpf | 11dpf |
| WT x WT 1 | 163;-;- | 159;0;4 | 159;0;4 | 159;0;4 | 159;0;4 | 159;0;4 | 159;0;4 | 159;0;4 | 159;0;4 | 159;0;4 | 159;0;4 |
| WT x WT 2 | 132;-;- | 132;0;0 | 131;0;1 | 131;0;1 | 131;0;1 | 131;0;1 | 131;0;1 | 131;0;1 | 131;0;1 | 131;0;1 | 131;0;1 |
| WT x WT 3 | 73;-;- | 73;0;0 | 73;0;0 | 73;0;0 | 73;0;0 | 73;0;0 | 73;0;0 | 73;0;0 | 73;0;0 | 73;0;0 | 73;0;0 |
| *vcc42/vcc42* x *vcc42/vcc42* 1 | 137;-;- | 117;6;14 | 107;15;15 | 107;6;24 | 107;2;28 | 107;2;28 | 107;0;30 | 106;0;30 | 106;0;30 | 106;0;30 | 106;0;30 |
| *vcc42/vcc42* x *vcc42/vcc42* 2 | 153;-;- | 143;2;8 | 137;8;8 | 136;6;11 | 135;3;15 | 134;0;19 | 134;0;19 | 134;0;19 | 134;0;19 | 134;0;19 | 134;0;19 |
| vcc42/vcc42 x vcc42/vcc42 3 | 102;-;- | 63;12;27 | 46;27;29 | 45;24;33 | 45;10;47 | 45;6;51 | 45;3;54 | 45;2;55 | 45;2;55 | 45;0;57 | 45;0;57 |
| *vcc42/vcc42* x *vcc42/vcc42* 4 | 265;-;- | 208;20;37 | 191;36;38 | 191;19;55 | 191;10;64 | 191;4;70 | 191;3;71 | 191;1;73 | 191;1;73 | 191;1;73 | 191;0;74 |
| Scramble controls | 33;-;- |  | 31;0;2 |  | 31;0;2 |  | 31;0;2 |  |  |  |  |
| *hmox1a* crispants | 32;-;- |  | 27;3;2 |  | 26;3;3 |  | 26;1;5 |  |  |  |  |
| *hmox1b* crispants | 81;-;- |  | 72;3;6 |  | 68;7;6 |  | 68;4;9 |  |  |  |  |
| *hmox1a+1b* crispants | 74;-;- |  | 57;14;3 |  | 56;2;16 |  | 56;1;17 |  |  |  |  |
| CuPP | 97;-;- |  | 96;0;1 |  | 95;0;2 |  | 95;0;2 |  | 95;0;2 |  |  |
| SnPP | 97;-;- |  | 95;0;2 |  | 93;0;4 |  | 90;0;7 |  | 83;0;14 |  |  |
| Scramble controls (WT) | 37;-;- |  | 37;0;0 |  | 36;1;0 |  | 36;1;0 |  |  |  |  |
| *hmox1b* crispants (WT) | 40;-;- |  | 38;2;0 |  | 38;1;1 |  | 38;1;1 |  |  |  |  |
| Scramble controls (*vcc42/vcc42*) | 74;-;- |  | 68;6;0 |  | 68;5;1 |  | 66;2;6 |  |  |  |  |
| *hmox1b* crispants (*vcc42/vcc42*) | 116;0;0 |  | 109;7;0 |  | 107;6;2 |  | 99;5;12 |  |  |  |  |
